# Selective attention network in naturalistic auditory scenes is object and scene specific

**DOI:** 10.1101/2025.01.03.631190

**Authors:** Patrik Wikman, Ilkka Muukkonen, Jaakko Kauramäki, Ville Laaksonen, Onnipekka Varis, Christopher Petkov, Josef Rauschecker

## Abstract

Everyday auditory scenes often contain overlapping sound objects, requiring selective attention to isolate relevant objects from irrelevant background objects. This study examined how selective attention shapes neural representations of naturalistic sound scenes in the auditory cortex (AC). Using functional magnetic resonance imaging, we recorded brain activity from participants (n = 20) as they attended to a designated object in scenes comprising three overlapping sounds. Scenes were constructed in two manners: one where each object belonged to a different category (speech, animal, instrument) and another where all objects were from the same category. Attending to speech consistently enhanced activations in lateral AC subfields, while attention to animal and instrument sounds preferentially modulated medial subfields, supporting models where attention modulates feature-selective neural gain in AC. Remarkably, however, spatial pattern analysis revealed that the attended object dominated the AC activation patterns of the entire scene in a manner depending on both object type and scene composition: When the objects of the scene belonged to different categories, attended objects dominated fields processing higher-level category-specific features. In contrast, when all scene objects shared the same category, dominance shifted to fields processing low-level acoustic features. Thus, attention seems to dynamically prioritize the features offering maximal contrast within a given context, emphasizing object-specific patterns in feature-similar scenes and category-level patterns in feature-diverse scenes. Our results support models where top-down signals not only modulate gain but also affect several steps of auditory scene decomposition and analysis – influencing stream segregation and gating of higher-level processing in a contextual manner, adapting to specific auditory environments.

## Introduction

We are surrounded by auditory objects, that is, sounds that can be assigned to a certain acoustic source, such as a person talking, someone playing a guitar by or a bird singing. The moment-to-moment auditory environment comprising such overlapping auditory objects form an auditory scene. Unlike in vision, the definition of what constitutes an auditory object is controversial (Kubovy and Van Valkenburg, 2001; Griffiths and Warren, 2004; Scott, 2005; Winkler et al., 2009; Bizley and Cohen, 2013). Nonetheless, it has been proposed that sound representations become increasingly abstract in the hierarchical auditory ventral “what” stream, proceeding from selectivity for simple acoustic features such as frequency in primary auditory regions, to complex acoustical structures (bandpass noise bursts, frequency-modulated sweeps) in secondary auditory regions, and culminating in full object representations in the anteroventral auditory cortex (AC) and categorical representations in middle temporal gyrus (Rauschecker et al., 1995; Rauschecker and Tian, 2000; Zatorre and Belin, 2001; Boemio et al., 2005; Petkov et al., 2008; Petkov et al., 2009; Rauschecker and Scott, 2009; Lewis et al., 2011; Rauschecker and Romanski, 2011; Theunissen and Elie, 2014; Rauschecker, 2018). There is still, however, considerable debate whether auditory objects are instead encoded through distributed representations within AC subfields selective for different low-level acoustic features (Formisano et al., 2008; Staeren et al., 2009; Leaver and Rauschecker, 2010b; Giordano et al., 2013; Giordano et al., 2023).

One important outstanding question is the role of selective attention on auditory object perception and their neural representation in AC (Bizley and Cohen, 2013). While both object formation and parsing of auditory scenes into their constituent streams seem to operate to some extent without attention (Sussman et al., 2002; Winkler et al., 2009), selective attention is thought to facilitate and transform both auditory object representations and auditory scene parsing (Zatorre et al., 1999; Cusack et al., 2004; Shamma et al., 2010). Especially, in natural scenes with several overlapping and competing auditory objects, attention is thought to play a pivotal role in forming stable object percepts (Shinn-Cunningham, 2008; Shamma et al., 2010; Shinn-Cunningham et al., 2017). Recently, studies on selective attention to “cocktail-party” speech has established that neural activity in AC reflects attended speech more accurately than the ignored speech (Mesgarani and Chang, 2012; Wikman et al., 2024b). However, due to the paucity of studies comparing the effects of selective attention on speech representations to that of other natural object categories, it is unclear whether this generalizes across auditory object categories (Hausfeld et al., 2018). Furthermore, it is still debated whether selective attention in natural scenes mainly operates on the feature level (Da Costa et al., 2013), object level (Zatorre et al., 1999; Shinn-Cunningham, 2008) or whether both are utilized in an adaptive contextual and task dependent manner (Shamma et al., 2010; Shamma et al., 2011; Wikman et al., 2020). Therefore, although selective attention can increase the gain of attended features at multiple levels, how auditory objects in naturalistic scenes are selected for remains unknown.

In the present functional magnetic resonance imaging (fMRI) our aim was to determine whether selective attention to different classes of natural sounds differentially modulates AC sound processing. To this end we presented natural sound clips, falling into three sound object categories (speech, animal, instrument), to participants (n = 20) in three different experiments (Fig 1). Participants selectively attended to one of three overlapping auditory objects or performed a visual control task. The three auditory objects were either from different sound categories *(three objects across, 3OA*) or from the same sound category (*three objects within, 3OW*). The experiments were designed to answer key questions on the role of selective attention in auditory scene analysis: (1) Does selective attention in naturalistic scenes operate primarily on feature or object-level representations? We conjectured attention may operate on low-level features only if they offer the necessary contrast to segregate the objects of the scene (Wikman et al., 2020). For instance, the level of attentional selection should depend on the sound scene, with low-level features being emphasised when scenes comprise objects from the same category of sound, with overlapping features, (*3OW*) in contrast to scenes with distinctly different objects, with few overlapping features (*3OA*). (2) What are the potential neural mechanisms underlying attentional selection of sound objects? Attention may simply increase the gain of AC neurons selective for auditory object stimulus properties (Woldorff et al., 1993; Hillyard et al., 1998; Zion-Golumbic et al., 2013). Alternatively, attention may also increase the selectivity of neural signals to objects and recruit higher order brain areas (Petkov et al., 2004; Rinne et al., 2012). Finally, we wanted to determine whether the attended object’s activation pattern dominated the activation pattern of the whole scene of overlapping auditory objects (Hausfeld et al., 2018; Hausfeld et al., 2021; Puschmann et al., 2024). Therefore, we conducted a third experiment (*Object-alone experiment, OA*) where participants listened to each of the auditory objects presented alone (to derive the activation pattern for each object type).

**Figure 1.**
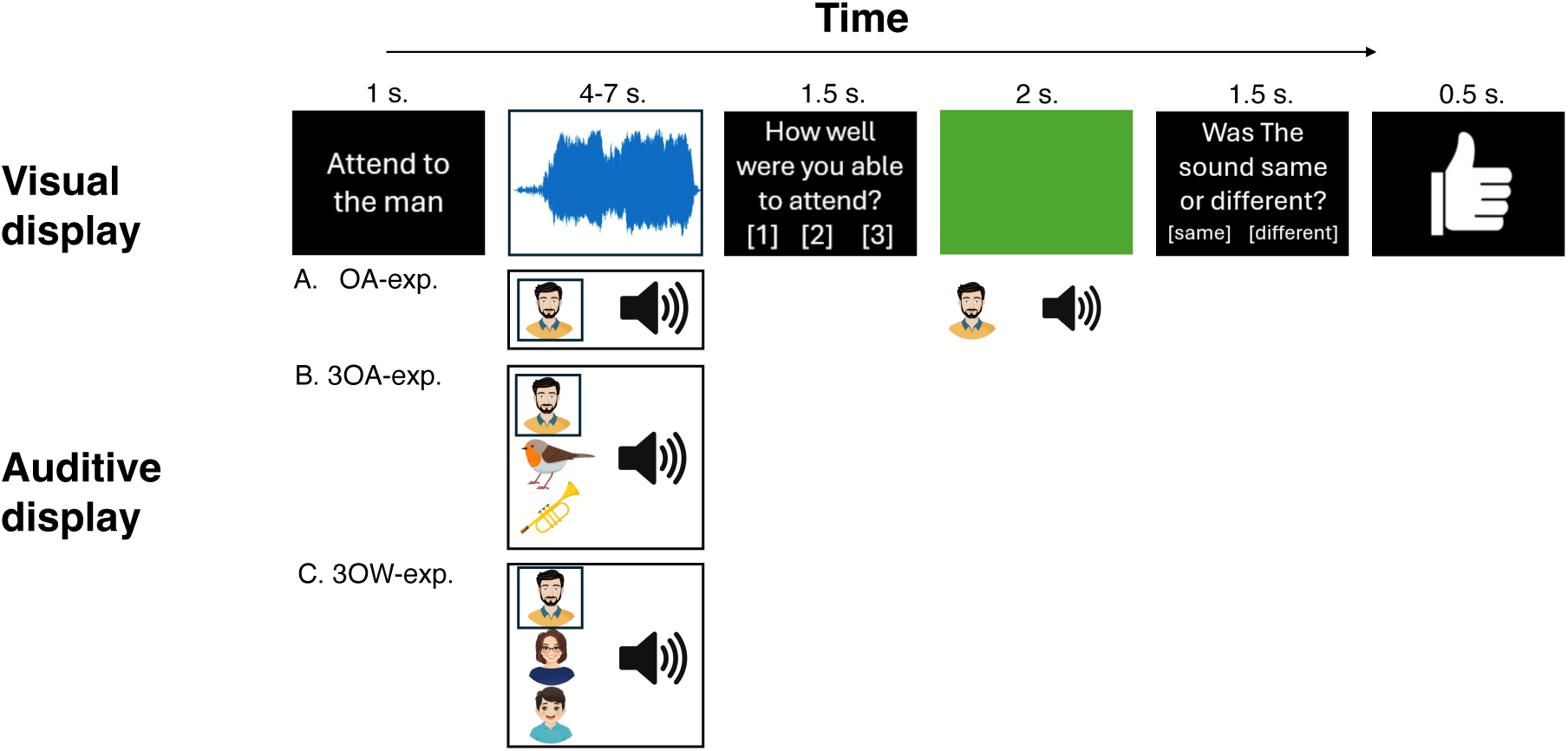
Behavioural paradigm used in the three experiments (*OA, 3OA, 3OW*). The trial structure was the same for all experiments, tasks and stimulus types. The trial started with an instruction (1s), orienting the participant to attend to a specific object. The attended object was either auditive (*attend sound object task*) or visual (*visual control task*). Thereafter the participant was presented with either one auditory object (OA), or three auditory objects (*3AO or 3OW*), as well as a visual video of a scrolling waveform display that lasted exactly for the time of the auditory stimulation. Participants attended either to the designated sound object (male speaker in the schematic illustration, “attend to man”) in the *attend sound object task* trials, while ignoring the video display; or to the video display while ignoring the auditory stimulation (“attend to visual display”, *visual control task).* In the *attend sound object task* after each stimulus trial, the participants rated their ability to attend to the designated auditory object and ignore the task irrelevant stimuli (1: less than 33% of the time, 2: 33–66% of the time and 3: more than 66% of the time). On some trials the participant also performed a match-to-sample task, where a short sample of the attended auditory clip was played or another clip from the same sound object. The participant was then tasked to report whether the sample was sampled from the previous attended object or from a different object. Lastly the participant receives feedback based on the response to the match-to-sample task. For details on the *visual control task* see *Trial structure*

## Materials and Methods

### Participants

Twenty healthy right-handed *monolingually* native Finnish speaking participants underwent an fMRI session (8 females, age range 19–32, mean age 22.7 years, standard deviation 3.68). All participants *were* adult university students at the University of Helsinki and Aalto University, had normal hearing, and normal or corrected-to-normal vision and no history of developmental or neurological disorders. We assessed handedness by using the Edinburgh Handedness Inventory *(Oldfield, 1971)* and acquired written consent from the participants before the study.

### Ethics Statement

For their participation, the participants were compensated monetarily 15 €/h. Each participant provided written informed consent. Written informed consent was obtained for the sharing of processed anomynised data from each participant. The experiment was accepted by the University of Helsinki Ethical Review Board in the Humanities and Social and Behavioural Sciences, and we conducted the study in accordance with the Declaration of Helsinki.

### Data availability statement

All data (behaviour and fMRI) pre-processed to anonymise, and original code have been deposited at Open Science Framework under Attention and Memory networks (HTTPS://DOI.ORG/10.17605/OSF.IO/AGXTH).

### Stimuli

The auditory stimuli comprised sound clips from three sound object categories: speech, animal, and instrument sounds. Each sound category included six subcategories. The speech subcategories comprised sound samples from two females, two males, and two children (one boy and one girl). The animal subcategories were sound samples from huskies, cats, whales, monkeys, birds, and seals. The instrument subcategories were sound samples from a baritone guitar, a 12-string guitar, a saxophone, a trumpet, an organ, and a bebop organ. Eight sound object clips (*exemplars*), from each subcategory, were used, making the total amount of different auditory clips 144 (3 categories (speech, animal, instrument), 6 sound objects for each category, and 8 exemplars from each sound object). The average duration of a sound sample was 5.3 s (4–7 s).

Speech samples were extracted from spoken dialogues recorded for our previous studies (Leminen et al., 2020b; Ylinen et al., 2024), where details are given about the recordings. The dialogues were related to emotionally neutral everyday situations. The speech samples were segmented into 4–7 second audio clips with Audacity (version 3.3.3), each comprising one sentence.

Animal sounds were collected from free internet sources. The sounds were selected based on the versatile range of produced sounds: for example, a howling husky with changing pitch was preferred over a monotonously barking dog. Soundtracks with as little background noise as possible were adopted for further processing. Selected sounds were transferred to Logic Pro X Waves X-Noise, where the remaining background noise was filtered with Channel EQ by dampening the noise by 20 – 50 dB depending on the sample.

Instrumental sound samples were created using clips from the same speakers as those used for the speech sounds, separate speakers being used for each specific instrument. Specifically, male voices were paired with 12-string guitar bright and baritone guitar samples, female voices with alto saxophone and trombone samples, and child voices with bebop organ and grand organ samples. The selection of instruments corresponding to each speaker was determined by the frequency characteristics of their voice, ensuring the best match between voice and instrument. For instance, a baritone guitar was chosen to correspond with a low male voice, as its natural frequency range aligned well with that of the voice. Initially, the frequency and duration of each dialogue was analysed using the Melodyne plug-in within Logic Pro X. Subsequently, the frequencies were adjusted to conform to the 12-Tone Equal Temperament (12-TET) scale, and MIDI files were generated for each dialogue. To ensure coherence, outlier frequencies in the original dialogues were adjusted to align with the general frequency range of each dialogue. Following this, the MIDI files underwent compression to ensure consistent volume (velocity in Logic Pro X) and timbre across notes. Additionally, an Audacity compressor was applied to further compress the MIDI files, with parameters set to a threshold level of – 30 dB, noise floor of – 35 dB, ratio of 5:1, attack time of 1.11 sec., and release time of 1.0 sec. The instrument samples utilized were predominantly original samples from Logic Pro X, except for organ samples sourced from the Logic Vintage B3 Organ plug-in.

All auditory stimuli were of high quality (48 kHz sampling rate and 16 bit). If the sounds had excessive internal variability, they were compressed with the Audacity compressor. After compressing, silent parts from the beginning and end were removed, and a slight Fade-in and Fade-out (50–100ms) was applied in Audacity. If there was noise in the audio clips, they were edited with the Auto-heal tool from Adobe Audition (version 22.6.0.66). The auditory stimuli were normalized using Matlab. Each audio file was scaled to adjust its average loudness to a target level measured in sones. The scaling was performed using the Matlab R2022b acousticLoudness() function, which returns loudness in sones according to ISO 532-1 (Zwicker) standard. The Matlab script determines the optimal scaling factor by minimizing the difference between the original and target loudness, computed as the average loudness across all audio files, using the fminsearch function. The audio signal was then scaled accordingly using the calculated factor. Only channel 1 was utilized in the computation. Finally, we filtered all sounds using the filter bank provided by Sensimetrics (model S14; Sensimetrics, Malden, Massachusetts, USA), to make the sounds optimal for the headphones used during fMRI (see *Equipment and Stimulus delivery*). At least 12 sound object exemplars were initially generated, of from which 8 were used in the actual trials. Out of the remaining 4 samples one was used for the practice and other 3 for the match-to-sample task.

The visual videos were created from all the auditory stimuli used in the experiment (214 sounds including the training sounds and sounds used as non-targets in match-to-target task). Using Matlab (R2020a, MathWorks Inc., Natick, MA, USA), for each sound a scrolling waveform display was generated with one-second window visible (blue waveform on white background, single audio channel normalized to maximize display size, concatenated twice and audio padded by a second of silence to have a smooth ending). Audio concatenation was done so that the generated videos would not end before the auditory stimulus presentation: video duration always outlasted the auditory stimulus during simultaneous presentation. The single waveform display frames were cropped to 876×656 pixel dimensions. From these frames, 25 FPS silent videos were created in AVI container format using ffmpeg and MPEG4 video codec with 2000 kbps target bit rate; AVI was used for compatibility issues. The resulting videos were manually screened to remove visually too similar videos, resulting altogether 180 different control videos used in the experiment, presented independently of the auditory stimulus.

### Stimulus acoustic features

Acoustic features were calculated with MIRtoolbox (v 1.8.1.) running in Matlab (2023b). Six features were calculated: *pitch*, *centroid*, *harmonic-to-noise-ratio* (*HNR*), *amplitude modulation standard deviation* (*AMSD*), *frequency modulation standard deviation* (*FCSD*), and *entropy*. For all features, a single value was derived for each clip. Pitch was calculated with the default autocorrelation method of the *mirpitch-*function, selecting the strongest periodic component (limited between 75 – 10000 Hz). Centroid (*mircentroid*) is the spectral centroid, the mean of the spectrum of the audio. HNR is calculated as the ratio of the strongest periodic component (highest autocorrelation) to the aperiodic component (Boersma and Weenink, 2001; Leaver and Rauschecker, 2010b). AMSD represents the variation in the sound amplitude across time and was calculated as the standard deviation of the absolute value of the sound signal after resampling to 60Hz. FCSD is the spectral variation in the stimulus, calculated as the standard deviation of power across frequency spectrum. Entropy is the relative Shannon entropy of the spectrum of the stimulus (*mirentropy*).

### Trial structure

Participants underwent three separate experiments (*object alone, OA; three objects across, 3OA; three objects within, 3OW*). The main differences between the experiments were that *OA* included only one sound object per trial, while *3OA* and *3OW* had sound scenes comprising three overlapping sound objects, which were chosen across the three categories of sounds (speech, animal, instruments) in *3OA* and within the three object categories in *3OW*. All three experiments (*OA, 3OA, 3OW*) used the same core trial structure, which can be divided into four parts: (1) an instruction was presented indicating whether the participant performed an auditory task (*attend sound object task*) and the identity of the sound object the participant was to attend to (e.g., “attend to the bird”), or whether the participant performed a visual task (*visual control task*, “attend to the visual display”). The instruction was displayed for 1 second. (2) After the instruction, the participant was presented with one sound object in *OA* (e.g., bird or male speaker), and three simultaneous sound objects in *3OA and 3OW*. In addition, a randomly sampled (without replacement) visual video (4 –7 sec.) was presented for the same duration as the auditory sounds. (3) Following the stimulus presentation, the participant was asked to give a subjective rating on how well they had been able to attend to the designated object by pressing a button (1.5 sec. response window) with an appropriate option: 1: less than 33% of the time, 2: 33–66% of the time and 3: more than 66% of the time. (4) Lastly, in 25% of *attend sound object task* trials, the participant was asked to complete a match-to-sample task. In the *attend sound object* match-to-sample task, a short sample (2 sec.) of either the sound object exemplar they had been attending to was presented (target trial, 50% probability) or another exemplar of the same sound object (nontarget trial, clipped from one of the 4 sound object exemplars that were not used as attended stimuli), was presented (1.5 sec. response window). To alert the participant of the match-to-sample task, the background of the visual display was changed to green before (1s) and during the presentation of the sample. After answering, the participant received feedback on their performance (0.5 sec.; correct/incorrect). The *visual control task* match-to-sample task was otherwise identical to the auditory version, except instead of auditory clips, the participant was presented with a short sample (2 sec.) of either the video clip they had been attending to was presented (target trial, 50% probability) or another video clip (nontarget trial), was presented (1.5 sec. response window). To ensure participants did attend to the visual display in all *visual control tasks* the match-to-sample task was presented in 100% of trials instead of 25%. The intertrial interval was randomized to last from 201 to 5500 msec.

### Experiments

In the *OA* experiment, the participants performed only one type of task: In the *attend sound object task* participants attended to the presented auditory object, which was always presented alone without any other sound objects, whereafter they gave a subjective rating of how well they could focus on the sound and performed a match-to-sample tasks (see *Trial Structure* for details). Note that to make the stimulus presentation comparable across the three experiments, visual videos were also present in the *OA*-experiment, although, participants were never instructed to pay attention to them in the experiment. For each participant each sound object exemplar (altogether 144) sound objects were presented in a separate trial. However, the presentation was split into four consecutive runs (36 exemplars/run) and each run always comprised two trials of each sound object type. The order of trials was randomly generated for each participant. For each sound object type there was one match-to-sample task where the sample was the same as the attended target sound object one where it was not. As the match-to-sample task was presented in 25% of the exemplars in one run, there were nine auditory match-to-sample tasks in each run.

In the *3OA-*experiment, unlike the *OA-experiment*, a *sound object scene* consisting of three sound objects, each from different sound categories (speech, animal, instrument) were presented to the participant (e.g., boy, husky, and trumpet). The participant performed two tasks (*attend sound object task* or *visual control task).* In this experiment during the *attend sound object task* the non-attended sound objects acted as distractors for the designated sound object that was to be attended. Accordingly, the participants were instructed prior to the experiment to not only attend to the designated sound object, but to also actively block out all distractors (auditory and visual). To help the participant with orienting to the desugnated sound object its presentation started 250-500ms prior to the distractor sound objects. In the *visual control task* participants were instructed to fully ignore all auditory stimulation and attend to the visual video. The attend sound object task was performed 75% of the trials and the rest were *visual control task* trials.

In the *3OA-*experiment sound objects for each trial were combined accordingly: All sounds were selected between sound object categories (speech, animal, instrument) and randomized individually for every subject. Every sound object type (e.g., *Male1*) was presented in two different auditory scenes during the *attend sound object task* (e.g. see A for each subcategory in Table 1). Every sound type acted twice as a target, once in both auditory scenes. Even though the same sound objects were presented together multiple times, the exemplar was always a different one so that in each run 108 exemplars were used only once during the *attend sound object task* (12 different types of scenes x 3 sound objects per scene x 3 different attended object conditions). The combinations were generated by arranging sound objects into three rows stacked vertically and then rotating the row of instruments one step and the row of animals two steps (for demonstration see bolded parts in Table 1). Because there were three possible task allocations of each sound object (attended, distractor 1, distractor 2) and two possible scenes for each of the six subcategories, the number of trials of *attend sound object task* were 36 in each run (3 attended objects x 12 sound scene types). To have a control for each sound object scene, we additionally generated *visual control task* sound object scenes each with the same sound objects as in the *attend sound object task* but different exemplars for each scene. Thus, we used the remaining 36 sound object exemplars for the *visual control task* (12 different types of scenes x 3 sound objects per scene). The exact same auditory scenes were presented in every run, but their order was randomized for every run.

**Table 1.**
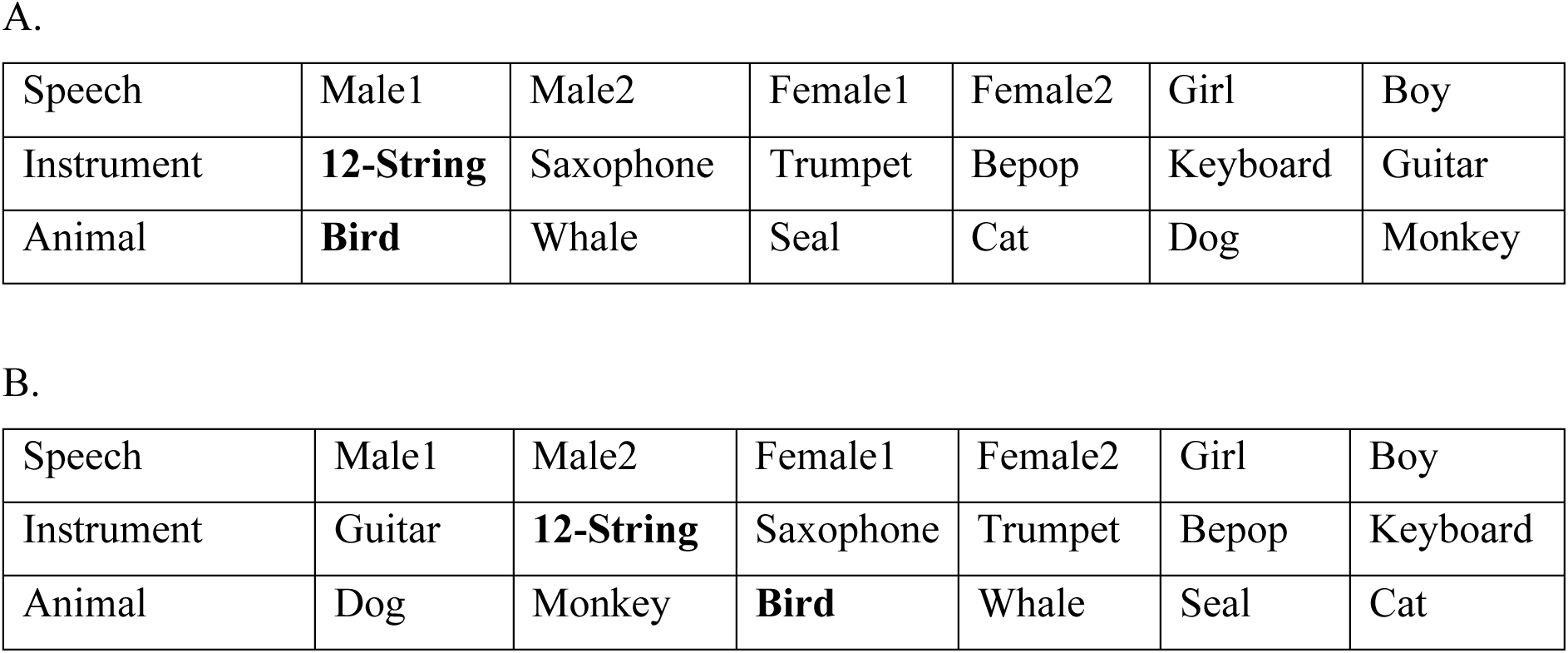
An example of a randomized stimulus structure of all possible 3OA-experiment scenes for one subject: Every sound object type (e.g. Male1) was presented in two different scenes (A and B). For example, Male1 could be presented either with a 12-string guitar and a Bird (e.g. first scene in A) or with a Guitar and a Dog (e.g. first scene in B). A new combination from A to B is created by rotating the instrument row one step and the animal row two steps forward. Note that the resulting 12 scenes were presented (with individual exemplars) in three different attend sound object versions. That is, the first scene (Male1, 12-string, Bird) were presented so that either Male1 was the focus of attention or 12-String or Bird, respectively. All 12 sound scenes were also presented during the visual control task, with different exemplars than the one used for the auditory tasks.

|  |  |  |  |  |  |  |
| --- | --- | --- | --- | --- | --- | --- |
| Speech | Male1 | Male2 | Female1 | Female2 | Girl | Boy |
| Instrument | **12-String** | Saxophone | Trumpet | Bepop | Keyboard | Guitar |
| Animal | **Bird** | Whale | Seal | Cat | Dog | Monkey |
| B. |  |  |  |  |  |  |
| Speech | Male1 | Male2 | Female1 | Female2 | Girl | Boy |
| Instrument | Guitar | **12-String** | Saxophone | Trumpet | Bepop | Keyboard |
| Animal | Dog | Monkey | **Bird** | Whale | Seal | Cat |

The trial structure and tasks for the *3OW-*experiment, was the same as that of *3OA*. However, in this experiment three sound objects were always from the same category of sound objects (e.g., Male1, Boy, Female2 or Bird, Husky, Whale). As in the *3OA-experiment,* each sound object type was presented in two different auditory scenes. However, in the *3OW-experiment* the allocation of sound objects to their auditory scenes could not be generated in the same way as in the *3OA-experiment*. Thus, the combinations were generated within each sound category (e.g., animal) first by randomly shuffling the order of the sound object types. Thereafter the sound objects were chosen to a scene, in a manner so that each sound object was presented in two and only two auditory scenes. This resulted in four possible subsets for each participant (see Table 2). Thus, there were three possible task allocations of each sound object (attended, distractor 1, distractor 2) and four different sets of sound objects per sound object category and three different categories of sound objects resulting in 36 number of trials of the *attend sound object task* in each run (3 attended objects x 4 sound scene types x 3 sound object categories). Similarly, as in the *3OA-experiment* the 12 sound scene types were also presented during the *visual control task* but with different exemplars.

**Table 2.** An example of a randomized stimulus structure of all possible 3OW-experiment scenes for one participant and one of the sound object categories (Animals): Every sound object type (e.g. Monkey, see first row) was presented in two different scenes (see row 2-4). For example, Monkey could be presented either with a Whale and a Bird (column 1) or with a Dog and a Cat (column 2). Note that the resulting 4 scenes were presented (with individual exemplars) in three different attend sound object versions. That is, the first scene (column 1) was presented so that either Monkey was the focus of attention or Whale or Bird, respectively. All 12 sound scenes (3 sound object categories x 4 sound scene types) were also presented during the visual control task, with different exemplars than the one used for the auditory tasks.

| Sound objects | Monkey | Whale | Bird | Dog | Cat | Seal |
| --- | --- | --- | --- | --- | --- | --- |
|  | **Scene 1** | **Scene 2** | **Scene 3** | **Scene 4** |  |  |
|  | Monkey | Monkey | Whale | Bird |  |  |
|  | Whale | Dog | Dog | Cat |  |  |
|  | Bird | Cat | Seal | Seal |  |  |

### Experimental session

During fMRI, every participant started the session with the *OA-*experiment. Half of the participants completed the *3OA*-experiment second and the *3OW-*experiment third. The other half completed the *3OW-*experiment second and the *3OA-*experiment third. For each experiment 4 runs were completed, making 12 total experimental runs. Each run in the *OA-* experiment included 36 trials and lasted around 6 minutes. For the *3OA* and *3OW* each run consisted of 48 trials lasting around 8 minutes. In between runs the participant was given a small break and the opportunity to communicate with the researchers. After the experimental runs, anatomical images were collected from participants.

### Pre-trial

Before the fMRI session participants practiced the experiments outside of the scanner. Each experiment was practiced for a total of two runs. Initially, each experiment was practiced with more lenient response times. Subsequently, a second set of practice runs was completed at a pace consistent with the actual paradigm. To mitigate potential learning effects, the practice runs used of stimuli that were left out from the actual fMRI-experiments.

### Equipment and Stimulus delivery

Stimuli were presented with Presentation 24.0 software (Neurobehavioral systems, Berkeley, California, USA). The auditory stimuli were presented binaurally through earphones (Sensimetrics model S14; Sensimetrics, Malden, Massachusetts, USA). To minimize the head movement and to reduce the volume caused by the fMRI scanner a piece of foam padding was placed over participants’ ears. Auditory volume was adjusted individually for each participant before the session at ca. 85 dB. Visual stimuli were projected onto a mirror attached to the head coil. The participant answered the questions with an fMRI-compatible response pad with either the index-, middle- or ring finger of their right hand which was placed on a foam padding.

### fMRI data acquisition

We collected functional imaging data with 3T MAGNETOM Skyra whole-body scanner (Siemens Healthcare, Erlangen, Germany) using a 32-channel head coil, with 2 coils removed to make the stimuli visually accessible. We used T2* echo planar imaging which collected 45 continuous oblique axial slices (TR 1050 msec., TE 30 msec., flip angle 75°, field of view 21 cm, slice thickness 2.7 mm, in-plane resolution 2.7 mm × 2.7 mm × 2.7 mm). Each session consisted of 12 runs. For the first 4 runs (*object alone* experiment) we collected approximately 360 volumes per run, whereas for the last 8 runs (*3OA/3OW-*experiments) approximately 570 volumes were collected per run. After the functional imaging we collected a high-resolution T1 anatomical image (TE 3.3 msec., TR 2530 msec., voxel matrix 256 × 256, in-plane resolution 1 mm × 1 mm × 1 mm) for coregistration. Each fMRI session lasted approximately 100 minutes.

### Analysis of Behavioural data and Stimulus features

We used *JASP* (0.18.3, https://jasp-stats.org/download/) to analyse behavioural data from participants. All misses were interpreted as incorrect in the match-to-sample task and removed from the self-rating. The mean task performance (in both subjective ratings and match-to-sample) and standard error of mean was used to establish that the participants were performing the task as expected. To analyse participants’ performance during the *attend speech object task* across experiments, we performed a repeated-measures analyses of variance (ANOVAs) separately for the rating and match-to-sample tasks. The ANOVAs included two factors: Experiment (*OA, 3OA, 3OW*), Sound object category (*speech, animal, instrument*). The performance during the *visual control task* was analysed with repeated measures ANOVAS with the factor Experiment (*OA, 3OA, 3OW*). We report η_p_^2^, and Greenhouse-Geisser adjusted *df*s when sphericity was violated.

For each stimulus acoustic feature (see *Stimulus acoustic features*) we used *JASP* to analyse whether sound object categories differed in their acoustic feature (each sound object exemplar acted as a data-point). Repeated measures ANOVA was used with the factor Sound object category (*speech, animal, instrument*), We report η_p_^2^, and Greenhouse-Geisser adjusted *df*s when sphericity was violated.

### Preprocessing of fMRI data

Preprocessing of the fMRI data were performed using FEAT (FMRI Expert Analysis Tool) Version 6.00, part of FSL (FMRIB’s Software Library, http://www.fmrib.ox.ac.uk/fsl). Registration of fMRI volumes to the high-resolution structural image of the participant was carried out using FLIRT (Jenkinson & Smith, 2001; Jenkinson et al., 2002), and preprocessing included motion correction using MCFLIRT (Jenkinson et al., 2002), slice-timing correction, non-brain removal with BET (Smith, 2002), and high-pass temporal filtering (with a cut-off of 100 Hz). For all further fMRI analyses except the spatial pattern analysis (see *Spatial Pattern analysis*), the data were then projected to the Freesurfer (Fischl, 2012) average surface space (fsaverage) using the Freesurfer function mri_vol2surfs.

To generate confound regressors for the first-level analyses we used *fMRIPrep* 20.2.5 (Esteban, Markiewicz, Blair, et al., 2018; Esteban, Markiewicz, Goncalves, et al., 2018). Confounding time-series for framewise displacement (FD), were calculated based on the preprocessed BOLD. FD was computed using two formulations following Power (absolute sum of relative motions, Power et al., 2014) and Jenkinson (relative root mean square displacement between affines, Jenkinson et al., 2002). FD was calculated for each functional run, both using their implementations in Nipype (following the definitions by Power et al., 2014). A global signal confound time series was extracted within the whole-brain masks Additionally, a set of confounding regressors (the four first accounting for the majority of variance) were extracted to allow for component-based noise correction (aCompCor, Behzadi et al., 2007). Gridded (volumetric) resamplings were performed in a single interpolation step using antsApplyTransforms (ANTs), configured with Lanczos interpolation to minimize the smoothing effects of other kernels (Lanczos, 1964). Finally, 6 motion correction parameter timeseries were added as confound regressors.

### First- and second-level analysis of fMRI data

In the first-level analyses, a general linear model (GLM) was fit to the time series data of each voxel in each run, using FEAT (3 columns format). A separate GLM was run for the two sound scene experiments (*3OA and 3OW).* All GLMs included three regressors (one for each sound object category) for the timepoints when performed the *attend sound object task.* The attended sound object delineated the sound object category. For example, if the participants attended to the *Husky* sound object in a scene comprising *Husky, Male1 and Trumpet,* the timepoints were include in the *attend sound object task animal* regressor. We also included nuisance regressors for all match-to-sample task timepoints. We also included one for the *visual control task* in *3OA,* and three regressors (one for each sound object category) in the *3OW*.

In *3OW*, to estimate differences in stimulus dependent processing of the respective sound scene types (*speech, animal, instrument*), we defined pairwise contrasts between the three different sound scenes presented during the *visual control task*, during which participants did not pay attention to the sounds. Thereafter, we defined interaction effects for ANOVAs with the factors Experiment (*3OA, 3OW*), Task (*attend sound object, visual control*), separately for each sound object category. These interaction effects are conceptually the same as a pairwise comparison between ARMs in the 3OA-experiment and the 3OW-experiment. Finally, we wanted to estimate differences in ARMs between the different attended object category types (e.g., which regions show stronger ARMs for speech than animal sounds). In *3OA,* we simply defined the pairwise contrasts between the three different categories of the *attend sound object task* (*attend to speech, attend to animal, attend to instrument*). However, in *3OW* both the stimuli and the focus of attention differed between the three different *attend sound object tasks.* Thus, to yield comparable results to those from *3OA* the same pairwise comparisons were performed after first contrasting each *attend sound object task* category with its respective *visual control task.* All pairwise comparison contrast maps were averaged over the four runs.

### Group-level univariate analysis of fMRI data

Group-level univariate statistics were based on a two level-procedure using a one-sample t-test performed using the glm-fit function of the Freesurfer software. Clusters were defined using permutation inference (a robust method for controlling false discoveries; Greve and Fischl, 2009) in Freesurfer, with the initial cluster forming threshold z = 3.1, cluster probability p < 0.01 for the univariate analyses.

### Effects of acoustic features on attention related modulations

For the acoustic features (see *Acoustic features*), where the three sound object categories differed, we ran fMRI regression analyses. This was conducted so that for each attended object we extracted its demeaned acoustic feature value. Thereafter we ran fMRI regression analyses with one regressor of interest, where each value was the demeaned acoustic feature for the attended sound object in the scene, separately for each acoustic feature. Each GLM included the same confounds as our general univariate analyses (see *First- and second-level analysis of fMRI data*)

### Spatial pattern analysis

We further tested whether selective attention in the *3OW* and *3OA* experiments elicits similar neural patterns as when the attended object was presented in isolation in the *OA*-experiment. We ran a searchlight (radius 6mm) separately for each subject and each trial in *3OW/3OA*, and, for each searchlight-voxel, regressed the spatial pattern in *3OW/3OA* with the spatial patterns from *OA* for the corresponding stimuli, and tested whether the *OA-*pattern of the attended stimulus better explained the pattern in *3OW/3OA* compared to the *OA-*patterns of the distractor stimuli.

Specifically, for a given voxel *V* and trial *T* in *3OW/3OA*, the spatial pattern *P* related to that trial was taken by averaging volumes 3*^rd^* to 8*^th^* (from trial onset), and taking the spatial searchlight pattern around voxel *V*. The corresponding patterns for the isolated presentation of each of the three stimuli in trial *T* were taken from the *OA-*experiment, and used as regressors to gain separate estimates for how well the attended (*Att*) and distractor (*Dist1*, *Dist2*) stimuli explain the pattern *P.* This was repeated for each voxel, trial, and subject. We then calculated the differences between the beta-values for attended stimuli and the two unattended stimuli (*Att* vs. *Dist1*, *Att* vs. *Dist2*), and averaged the differences across trials and within category (*speech, instrument, animal*), experiment (*3OW* and *3OA*), and subject, resulting in wholebrain spatial pattern difference maps.

Furthermore, as in the *3OA*-experiment the distractors were always from different sound categories than the attended object, and these maps could thus show effects of object-category instead of selective attention. Therefore, we also subtracted from the mean category-specific *Att-*values (i.e., all *Att*s of that sound category) the mean of that same sound category’s values when they were distractors (all *Dist1*s and *Dist2*s of that category; *asDist1* and *asDist2*). Finally, to test whether category-level patterns (mean of the *OA*-patterns of e.g. all instruments) can explain the results, we added these as regressors, thus having 6 regressors (3 exemplars, 3 categories) in *3OA*, and 4 regressors (3 exemplars, 1 category) in *3OW*.

Group-level conjunction maps were performed in four steps steps: First, each spatial pattern difference map was projected to the Freesurfer average (fsaverage) using the participants’ own Freesurfer surface (surface smoothening: 5 *mm^2^* full width half maximum smoothening). Thereafter, for each spatial pattern difference map, we performed a one-sample t-test using the glm-fit function of the Freesurfer software, and defined clusters using permutation inference in Freesurfer, with the initial cluster forming threshold z = 2.3, cluster probability p < 0.05. Finally, in *3OW* we calculated the overlapping clusters for two spatial pattern difference maps (*Att vs Dist1* and *Att* vs *Dist2*), and in *3OA* we calculated the overlapping clusters for four spatial pattern difference maps (*Att vs Dist1*, *Att* vs *Dist2*, *Att* vs *asDist1*, and *Att* vs *asDist2*).

### Bayesian region of interest analysis

We used the significant clusters (initial cluster threshold, z = 3.1 permutated cluster significance, p < 0.01) to define our ROIs for Bayesian analyses. We calculated the mean Bold-signal for each condition/participant/ROI in the dataset that did not yield the significant cluster. Bayesian pairwise comparisons and ANOVAs were defined in *JASP*, and we report *bf_10, incl_* for the effects of interest.

## Results

The three experiments were designed to yield complementary information about differences in attention-related modulations (*ARMs*) between the three types of sound object categories. The *3OA* experiment was designed to reveal how selective attention modulates processing when attending to objects in scenes that comprise different categories of objects. In contrast, *3OW* was designed to reveal how selective attention modulates processing in scenes that comprise sound objects from the same object category. Furthermore, because in *3OW* each scene comprised of sound objects from only one category, we used data from this experiment to estimate how stimulus dependent processing differed between the three categories of sound (see below). The *OA*-experiment was used as a point of reference for behavioural performance (see *Behavioural performance*), as well as, in our spatial pattern analyses, to yield spatial activation patterns for single objects (see *Attention-driven modulation of spatial activation patterns*).

## Behavioural performance

To study task performance in the three experiments (*OA, 3OA, 3OW*) we used subjective ratings and match-to-sample tasks (see *Trial Structure*, and Fig 1). Our aim in the behavioural analyses was to evaluate whether participants successfully attended to the sound objects and whether attending to any of the object categories stood out in difficulty. To evaluate whether subjective ratings depended on the experiment and attended sound object category, we ran a repeated measures repeated measures ANOVAs with factors Experiment (*OA, 3OA, 3OW*) and Sound object (*speech, animal, instrument)* (Fig 2 A–C). There was a significant main effect of Experiment (F_2,38_ = 46.4, p < 0.001, η_p_² = 0.71), as participants found attending to the designated object easiest in *OA* (μ = 2.95, sem = 0.046), thereafter in *3OA* (μ = 2.71, sem = 0.046), and most difficult in *3OW* (μ = 2.45, sem = 0.046). There was a significant main effect of Sound object (F_1.5,38_ = 20.6, p < 0.001, η_p_² = 0.52), as participants rated speech and animal sound objects (μ = 2.78, sem = 0.041; μ = 2.75, sem = 0.041) easier to attend to than the instrument sound objects (μ = 2.57, sem = 0.041). There was also a significant Experiment x Sound object interaction (F_2.4,38_ = 8.68, p < 0.001, η_p_² = 0.314), because participants found it more difficult to focus their attention on the instrument sound objects than the other two sound object categories in the 3OA-experiment than the other two experiments (see Fig 2A–C).

**Figure 2.**
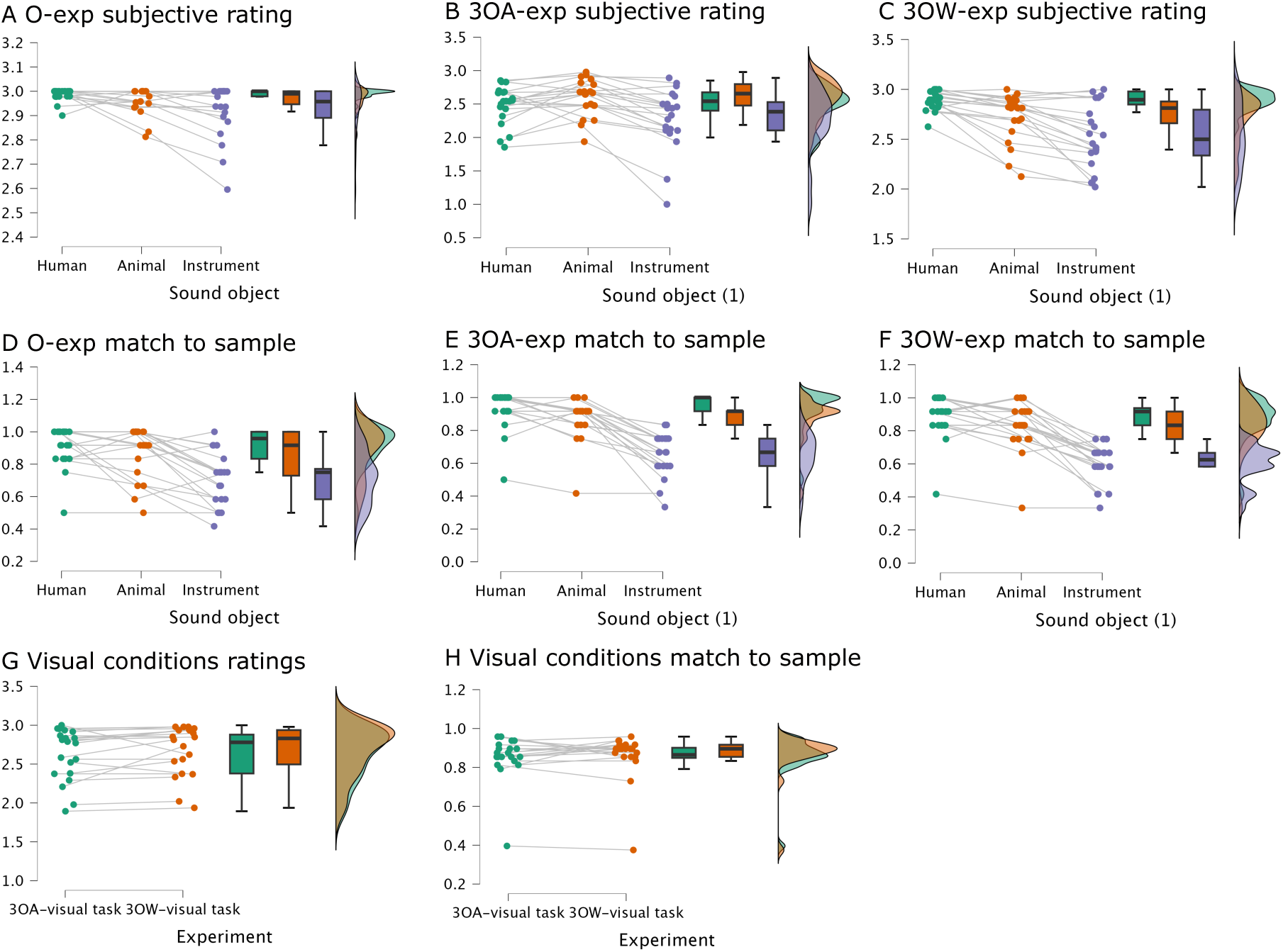
Behavioral results from the three experiments (*OA, 3OA, 3OW*). Participants gave both subjective ratings about their ability to attend to the stimuli and performed a match-to-sample task. In the rating task participants were instructed to rate how well they could focus on the attended stimuli (designated sound object in the auditory conditions and visual video in the visual control conditions) with three different options (1: less than 33% of the time, 2: 33–66% of the time and 3: more than 66% of the time). In the match-to-sample task participants were instructed to indicate whether a short 2s excerpt matched the attended object or not. Each subplot first depicts the participants’ behaviour across conditions (each line corresponds to a separate participant), thereafter box plots for each condition (median, ±25 percentiles and ±47.5 percentiles), and thereafter distributions for each condition. ***A-C*** Shows subjective ratings for each sound object type separately for the three experiments. ***D-F*** Shows performance in the match-to-sample task for each sound object type separately for the three experiments. ***G-H*** Shows subjective ratings, and performance in the match-to-sample task during the visual control conditions separately for the 3OA and 3OW-experiment respectively.

To evaluate whether participants performance in the match-to-sample tasks depended on the experiment and the sound object category we ran a repeated measures ANOVAs with the same factors as for the ratings (see Fig 2 D–F). There was a significant main effect of Experiment (F_2,38_ = 4.3, p < 0.05, η_p_² = 0.19), as participants performed worse in *3OW* (μ = 0.76, sem = 0.025), than *OA* and *3OA* (μ = 0.83, sem = 0.025; μ = 0.82, sem = 0.025). There was also a significant main effect of Sound object (F_1.4,38_ = 95.3, p < 0.001, η_p_² = 0.83), as participants performed better with speech and animal sound objects (μ = 0.82, sem = 0.025; μ = 0.81, sem = 0.025) than the instrument sound objects (μ = 0.75, sem = 0.025). However, there was no significant Experiment x Sound object interaction (F_3.2,38_ = 1.36, p > 0.05, η_p_² = 0.067).

To evaluate whether the behaviour in the visual control task depended on experiment, we ran for both subjective ratings and match-to-sample tasks a repeated measures ANOVAs with the experiment (*3OA, 3OW*) as a factor. These ANOVAs revealed that there were no differences in subjective ratings (F_1,19_ = 2.32, p > 0.05, η_p_² = 0.11), nor match-to-sample performance depending on the Experiment (F_1,19_ = 0.28, p > 0.05, η_p_² = 0.015) see Fig 2 G–F.

### Stimulus-dependent processing of sound scenes in AC

First, we wanted to determine whether there were significant differences in the processing of stimulus features between the three categories of sound objects (*speech, animal, instrument*) in AC, when the objects were not the focus of attention. To examine such stimulus-dependent processing, we conducted pairwise comparisons between the three different auditory scene types (scenes comprising only speech/animal or instrument sounds) presented during the *visual control task* in *3OW*, where participants focused on the visual stimuli and ignored the auditory scenes. In these comparisons all significant differences between the sound scenes were confined to the AC (for the definition and parcellation used for AC subfields see Fig. 5 and (Glasser et al., 2016)). Speech sound scenes were associated with significant stimulus-dependent processing in all STG/S subfields, as well as the subfields A4 and A5, when comparing to the other two sound object categories (Fig 3A–B, yellow and red). In contrast, animal and instrument sound scenes activated the AC subfields MBelt, 52, and PI more strongly than speech sound scenes (Fig 3A–B, blue and turquoise), while instrument sound scenes activated STGa, Ta2, and MBelt more strongly than animal sound scenes (Fig 3C, blue and turquoise).

**Figure 3.**
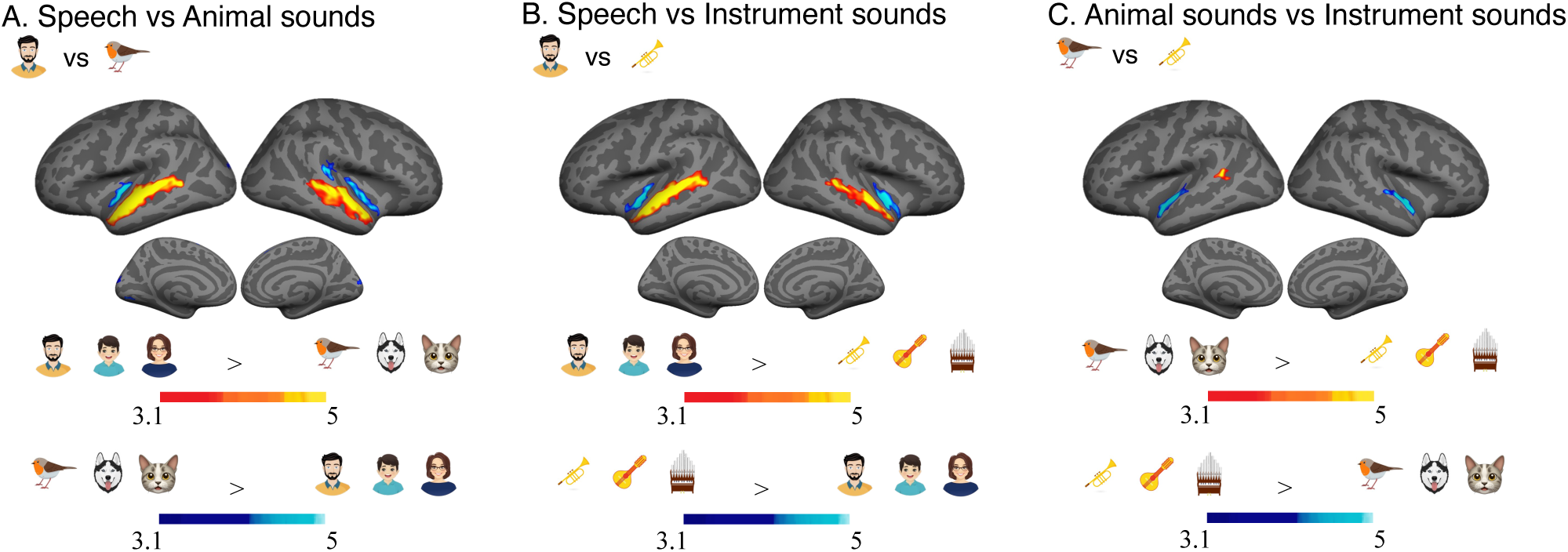
Stimulus-dependent processing of the three sound object category scenes (speech, animal, instrument) was estimated using pairwise comparisons between the three different auditory scenes presented during the *visual control task* in *3OW* (participants focused on the visual stimuli and ignored the auditory scenes). Group functional maps are shown overlaid on lateral and medial views of the freesurfer left and right inflated fsaverage surfaces (lighter gray denotes gyri and darker gray sulci). In all subplots, initial cluster threshold was z = 3.1; permutated cluster significance p < 0.01) ***A*** Shows differences in stimulus-dependent processing between speech (red and yellow) and animal (blue and turquoise) sound objects. ***B*** Shows differences in stimulus-dependent processing between speech (red and yellow) and instrument (blue and turquoise) sound objects. ***C*** Shows differences in stimulus-dependent processing between animal (red and yellow) and instrument (blue and turquoise) sound objects.

### Differences in attentional modulation of sound objects

Next, we delineated sound category selective *ARM* clusters separately for the two experiments (*3OA, 3OW*) using pairwise comparisons between the three respective object categories (*speech, animal, instrument*). In 3OA, *ARMs* can be estimated simply by performing pairwise comparisons between the respective conditions, since the auditory scenes always comprised three objects from three different object categories and only the focus of attention varied between the conditions. In contrast, in 3OW this is not the case, because the auditory scenes always comprised one category of objects, and thus, direct comparisons might be dominated by stimulus-dependent activity. Therefore, to yield comparable results to those from *3OA*, in *3OW* we extracted *ARMs* for each sound object category by first contrasting each *attend to sound object task* condition with its corresponding *visual control task* condition. Because each auditory scene (e.g., male speaker, child, female speaker) was presented both during the attend sound object task and visual control task, any stimulus-dependent processing should be factored out in these contrasts. Thereafter pairwise comparisons were performed similarly as for *3OA*. However, because in the *3OW* ARMs were extracted after first controlling for stimulus-dependent processing, it was to be expected that this experiment would have less power than the *3OA* (where conditions could be directly compared).

Pairwise comparisons between the three object category *ARMs* separately for the two experiments (*3OA, 3OW*) are displayed in Figure 4 (speech vs. animal sound objects, A, D; speech vs. instrument sound objects, B, E; animal vs. instrument sound objects, C, F). As can be observed by comparing the results displayed in Fig 3 with those in Fig 4, differences in stimulus-dependent processing between the three object categories were observed in Similar AC subfields as differences in *ARMs*. However, some differences between the sound object ARMs in AC were observed in subfields where no stimulus-dependent processing differences were observed: stronger *ARMs* were observed for human sound objects than animal sound objects in A1 (Fig 4A, red and yellow), and a cluster in the STSda showed stronger *ARMs* for animal than instrument sound objects (Fig 4C, red and yellow).

**Figure 4.**
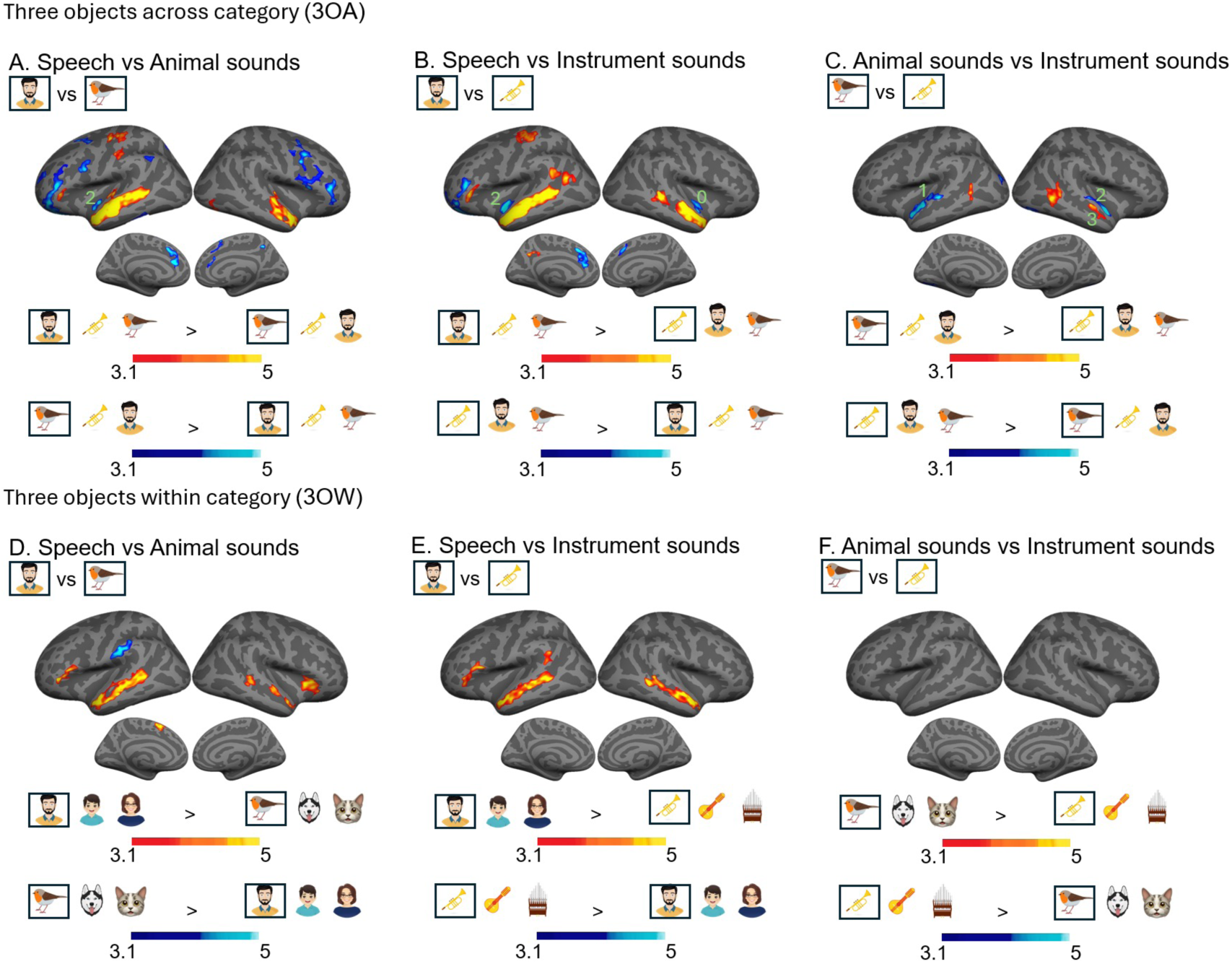
Pairwise comparisons reveal differences in attention-related activations between speech, animal and instrument sound objects. In all subplots we used an initial cluster threshold z = 3.1, permutated cluster significance p < 0.05, FWER. ***A, D*** Shows differences in ARMs between speech and animal sound objects in *3OA* and *3OW* respectively. ***B, E*** Shows differences in ARMs between speech and instrument sound objects in *3OA*, and *3OW* respectively. ***C, F*** Shows differences in ARMs between animal and instrument sound objects in *3OA* and *3OW* respectively. Note that generally differences between sound object ARMs are observed in overlapping AC sub-fields as stimulus-dependent processing (Fig 3). As some clusters in the AC were only observed in *3OA*, we ran a Bayesian pairwise comparisons for *3OW* data extracted from the cluster to test whether there was evidence for the same effect albeit subthreshold in *3OW*. In these Bayesian pairwise comparisons, we denote Bayes factor (bf_10_), 0: bf < 1, negative evidence; 1: 1 < bf < 3, weak evidence; 2: 3 < bf < 20, Positive evidence; 3: 20 < bf < 150, Strong positive evidence. Thus, the number 2 in ***A*** denotes that there is positive evidence that animal sound object related ARMs in this region are also present in the *3OW* dataset (***D***).

Outside the AC, significant differences between sound object *ARMs* were observed in inferior frontal and sensorimotor regions for speech objects (Fig 4 A–C, red and yellow), and various prefrontal, inferior parietal, medial frontal, and medial posterior subfields for the animal and instrument sound objects (Fig 4 A–C, blue and turquoise).

As was expected (see above), many of the smaller clusters observed in the *3OA* experiment did not survive cluster correction in the *3OW* experiment. Therefore, we extracted the significant AC clusters (that were not observed in the *3OW* experiment) of the *3OA*-experiment and ran a ROI-based Bayesian analysis (see *Bayesian region of interest analysis*) for the *3OW* data to test whether the *ARM* differed in these clusters between the experiments. As can be seen in Fig 4, there was generally positive–strong positive evidence (3 < bf_10_ < 20; 20 < bf_10_ < 150) that similar effects were present in the *3OW*-experiment (see Fig 4).

### Relationship between low-level acoustic features and sound object related attentional modulation in the auditory cortex

All sound object categories were matched by duration and perceptual loudness. In addition, speech and instrument sound object categories were rendered highly similar on other acoustic features, because the instrument sound object exemplars were generated from speech sound object exemplars of the same speakers as those that constituted our speech sound objects (see *Stimuli*). However, the animal sound object exemplars were gathered from heterogenous sources (see *Stimuli*) making it impossible to fully match acoustic features of the animal sound object category with the speech and instrument categories. Therefore, as in previous studies (Leaver and Rauschecker, 2010b), we assessed whether the categories differed on several acoustic features, including spectral (*pitch* and *frequency centroid*), spectral structure (*harmonic to noise ratio*), temporal variability (frequency modulation standard deviation and *amplitude modulation standard deviation*), and *entropy*. For this purpose, we used ANOVAs with the factor Sound object category (speech, animal, instrument) to test whether the object categories differed on any of these acoustic features. There were significant main effects of Sound object category for all studied acoustic features except *frequency modulation standard deviation* (see Fig7 middle column): *pitch* (F_2,85.3_ = 21.2, p < 0.001, η² = 0.28), *frequency centroid* (F_2,87_ = 16.8, p < 0.001, η² = 0.2), *harmonic to noise ratio* (F_2,89_ = 5.67, p < 0.01, η² = 0.05), *amplitude modulation standard deviation* (F_2,93.8_ = 22.89 p < 0.001, η² = 0.240), and *entropy* (F_2,81.7_ = 29.9 p < 0.001, η² = 0.190).

As the sound object categories differed on five of the six acoustic features studied, we examined the extent to which our category-selective *ARMs* (see Fig 5) were influenced by the fact that the categories differed in their acoustic features. For this purpose, we ran a separate regression analysis (see *Effects of acoustic features on attention related modulations*) for each of the five acoustic features on *3OA* fMRI data, which was chosen as *ARMs* were more strongly category specific in this experiment (see Fig 4). Note that the value for each acoustic feature was derived for the attended object of the sound object scenes (see *Effects of acoustic features on attention related modulations*). By comparing the results of these analyses (Fig 5.) with the category selective ARMs presented in Figure 4, it can be discerned that some of the clusters, such as, negative correlations for pitch and harmonic to noise ratio (Fig 5A,C, blue and turquoise), as well as, positive correlations for amplitude modulation standard deviation (Fig 5D red and yellow), arose in the same regions where speech sound objects differed from the two other object categories (Fig 4 A, D, B, E). Thus, one could argue that the human selective clusters displayed in Figure 4 arose because speech sounds differed on these acoustic features. However, because the same clusters arose for all the acoustic features, and the categorical effect (speech < animal, instrument; Fig 4) was much stronger than any acoustic feature effect, we argue for the reverse.

**Figure 5.**
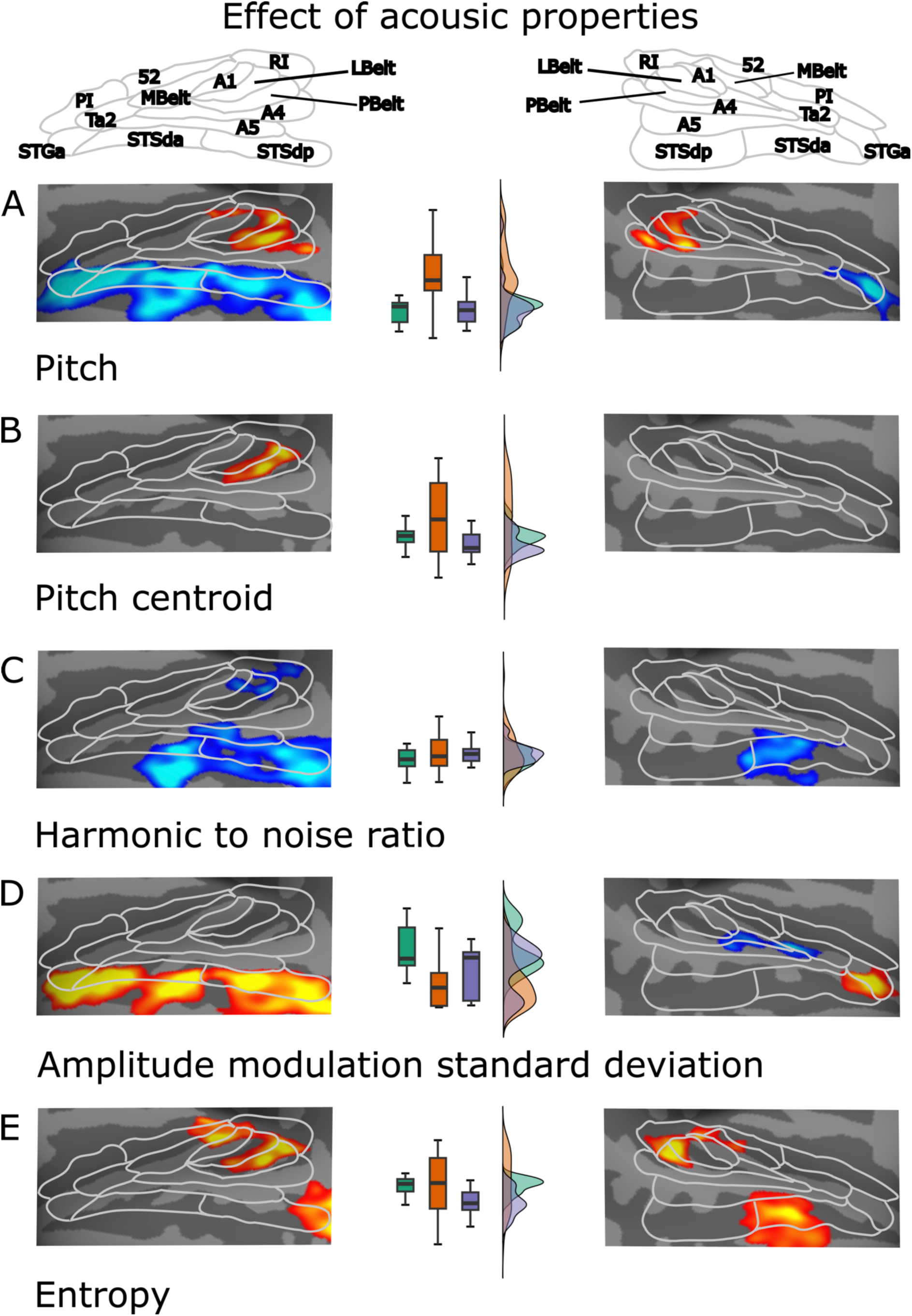
To examine the extent to which our category-selective *ARMs* were influenced by acoustic features of the attended sound objects, we performed separate regression analyses for each acoustic feature. Note that the value for each acoustic feature was derived for the attended object of the sound object scenes (see xxx). At the top of the figure, we show AC subfields as defined in the HCP-parcellation (Glasser et al., 2016). In each subplot (***A–E***), red and yellow denote significant positive correlations and blue and turquoise denotes significant negative correlations. In the middle of the subplots, we show box- and distribution plots of the respective auditory sound object category (*speech, animal, instrument*) acoustic feature values. In all subplots we used an initial cluster threshold z = 3.1, permutated cluster significance p < 0.01.

In contrast, the regression analyses for acoustic features also revealed correlations in AC subfields that were not observed for any ARM contrast between sound object categories. That is, pitch, pitch centroid and entropy correlated positively with *ARMs* bilaterally in LBelt and PBelt (Fig 5A,B,E, red and yellow); harmonic to noise ratio correlated negatively with *ARMs* in RI, LBelt and A1 (Fig 5C, blue and turquoise); amplitude modulation standard deviation correlated negatively with *ARMs* in A4 (Fig 5D, blue and turquoise); entropy correlated positively bilaterally in posterior parts of MBelt (Fig 5E, red and yellow).

### Differences between ARMs in *3OA* and *3OW*

We also wanted to determine whether ARMs differed when participants selectively attended to a sound object in auditory scenes comprising objects from the three different object categories (*3OA*) compared to when all objects were from the same object category (*3OW*). To this end, we inspected interaction effects from three separate (*speech, animal, instrument*) repeated measures ANOVAs with factors Experiment (*3OA*, *3OW*) and Task (*attend sound object task, visual control task*). This interaction effect is conceptually the same as a pairwise comparison between *ARMs* in the *3OA*-experiment and the *3OW*-experiment. As can be seen in Figure 6, *ARMs* were bilaterally stronger for *3OW* than *3OA* in the inferior frontal cortex for both speech and animal sound objects (Fig 6A–B), as well as anterior insula, and STG/STS regions for speech sound objects (Fig 6A). That is, most significant clusters were observed only for speech sound objects. However, in order not to make the wrong conclusion that these effects were exclusive to speech, we ran Bayesian ANOVAs to evaluate whether there was evidence for similar ARM differences for the two categories that did not yield significant clusters. Thus, we extracted the significant clusters and performed Bayesian ROI analyses for the two sound object categories that did not yield significant clusters (see *Bayesian ROI analyses*). In general, the clusters found for speech (and animals) yielded positive–strong positive evidence (3 < bf_10_ < 20; 20 < bf_10_ < 150) for other sound objects (see Fig 6).

**Figure 6.**
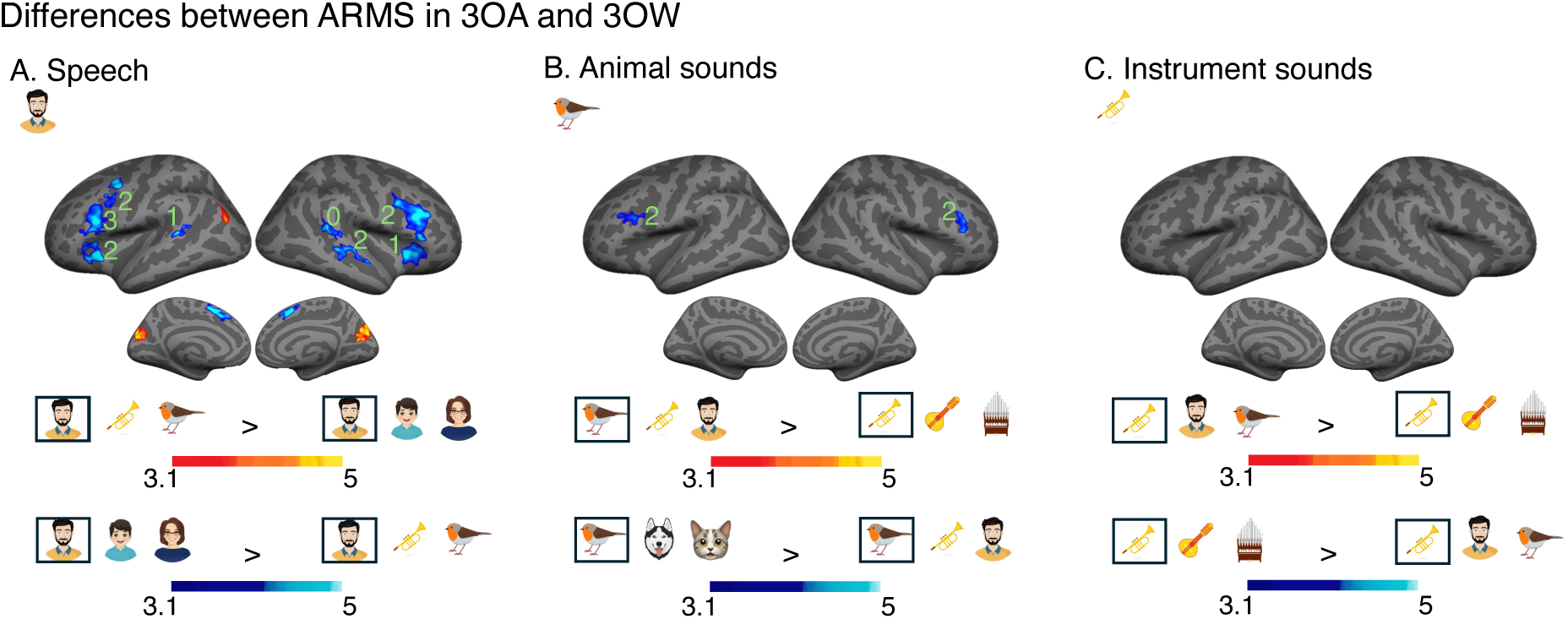
Repeated measures ANOVAs reveal differences between *ARMs* in the two experiments (*3OA > 3OW*, red and yellow; *3OW > 3OA*, blue and turquoise), performed separately for attended speech, animal or instrument sound objects. Note that from these ANOVAs we only present the interaction effects, as they are conceptually the same as pairwise comparisons of the estimated *ARMs* in the two experiments (see Fig 4). In all subplots we used an initial cluster threshold z = 3.1, permutated cluster significance p < 0.01. ***A*** Shows significant differences in *ARMs* for speech sound objects (red and yellow). ***B*** Shows significant differences in *ARMs* for animal sound objects. ***C*** Shows significant differences in *ARMs* for instrument sound objects. Note that in most of the clusters *ARMs* were stronger when the sound scenes comprised only one category of sound objects (*3OW*, blue and turquoise) than when the auditory scenes comprised all three categories (*3OA*, red and yellow). As some clusters were observed for only one of the three attended sound objects, we ran Bayesian repeated measures ANOVAs (Experiment (*3OA, 3OW*) and Sound object (object1, object2)), for data extracted from the cluster to test whether there was evidence for the same effect, albeit subthreshold. In these Bayesian analyses the same BF notation is used as in Fig 4. Thus, the number 3 in ***A*** denotes that there is strong positive evidence that ARMs are stronger in the *3OW* than the *3OA* experiment also for animal and instrument sound objects (***B, C***).

### Attention-driven modulation of spatial activation patterns

To evaluate whether the focus of attention not only affected mean BOLD signal but also spatial activation patterns, we determined whether the voxel patterns present in scenes comprising three separate objects (in *3OW* and *3OA*) were dominated by the voxel pattern of the attended object at the cost of the distractor objects. Thus, we derived voxel patterns (6mm radius searchlight) for each sound object exemplar from *OA-*experiment (where each object was presented in isolation) and used these voxel patterns as regressors to explain voxel patterns of scenes with the same sound object exemplars in the *3OW and 3OA-*experiment (for details on this analysis see *Spatial pattern analysis*). For example, in the *3OW*-experiment for a scene where exemplar no. 1 of the woman speech object was attended and exemplar no. 8 of the boy speech object and exemplar no. 4 of the male speech object were distractors, we derived the voxel pattern for each of the three object exemplars from the *OA*-experiment and used these voxel patterns as regressors for the voxel pattern of the full scene. To evaluate whether the attended object exemplars voxel pattern dominated the voxel pattern of the whole scene, we performed conjunction analyses for two pairwise comparisons in *3OW* and four pairwise comparisons in *3OA* between the correlation maps (see *Spatial Pattern Analysis* and Fig 7 for details).

**Figure 7.**
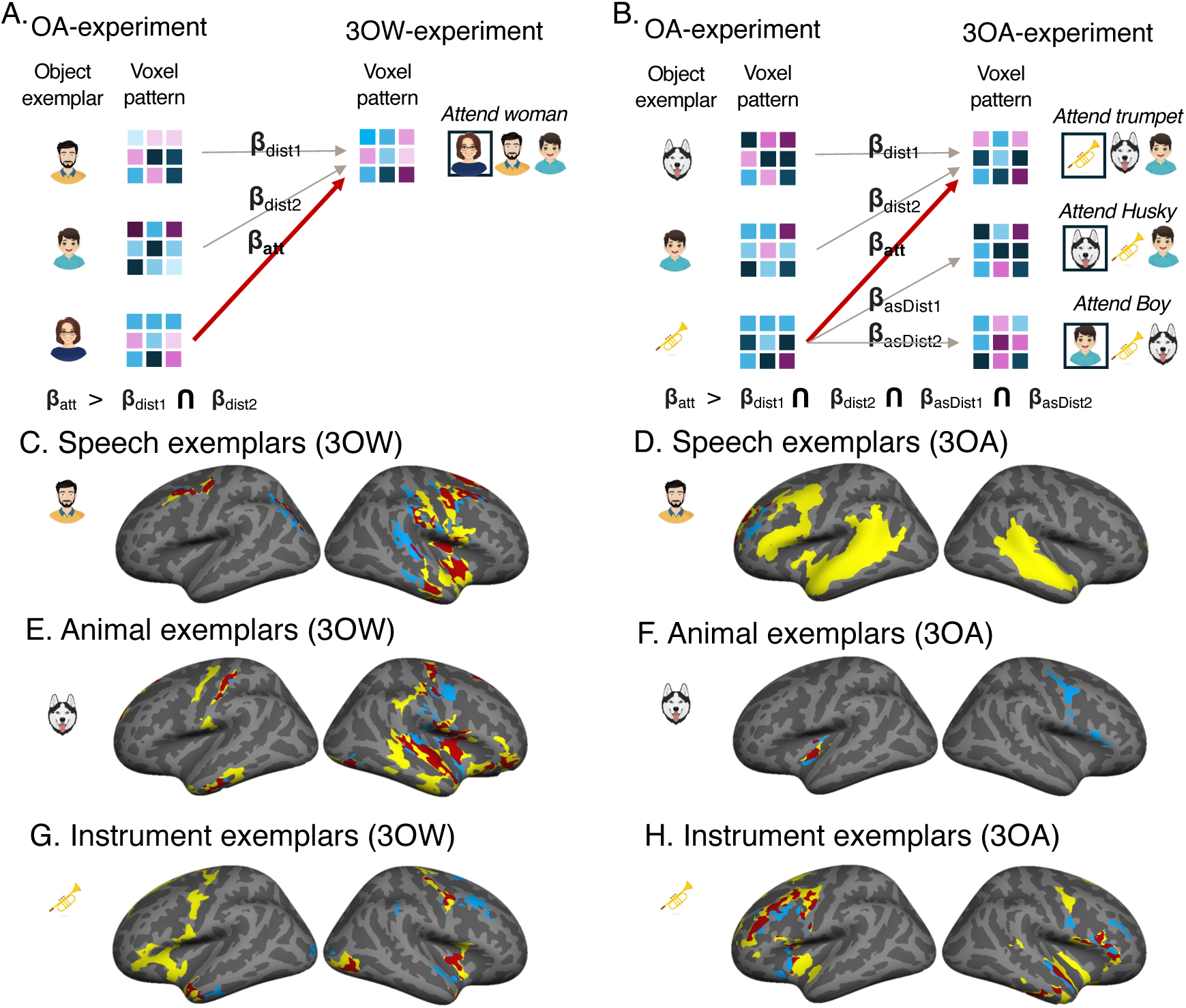
Results from combination analyses showing areas with higher correlations between spatial patterns across experiments for attended than unattended object exemplars separately for the three object categories (*speech, animal, instrument*). For both experiments (*3OW and 3OA*, ***A, B***) a searchlight analysis (6mm^3^ radius) was run, comparing spatial brain patterns elicited by the same specific auditory exemplar in the *OA* and the auditory scene experiments (*3OA and 3OW*). This was done separately for each stimulus exemplar and each subject, and correlations for each object category were averaged. ***A*** For *3OW* we used the voxel pattern of each of the three object exemplars present in the scene (in this illustration man, distractor1; boy, distractor2; woman, attended) from *OA* and regressed the voxel patterns with the voxel pattern of the full scene in *3OW*. This was done for each scene, whereafter we calculated average regression maps (**β**_att_, **β**_dist1_, **β**_dist2_) separately for auditory scenes where speech, animal or instrument objects were attended. Hereafter, we calculated two pairwise comparisons, for each object category separately (**β**_att_ > **β**_dist1_) and (**β**_att_ > **β**_dist2_) with an initial cluster threshold z = 2.3, permutated cluster significance p < 0.05, and show voxels where both comparisons yielded significant clusters. The result of these combination analysis is displayed separately for each object type in ***C, E, G***. Clusters are shown for two combination maps: red and yellow denotes clusters from the above-mentioned analyses, while red and blue from the same analysis, where all object category level (i.e. voxel pattern across all exemplars) has been controlled for in the regression analysis. I.e. the combination analysis shows for each object category the regions where the attended object voxel pattern significantly better predicted the voxel pattern of the scene than the two distractors. ***B*** For *3OA* the logic was the same as for *3OW*, with the exception that for each object category we also averaged the two instances when the same object exemplar was a distractor (**β**_asDist1_, **β**_asDist2_) and calculated pairwise comparisons between **β**_att_ and the **β**_asDist_ maps. ***D, F, H*** The result of the combination analysis was thereafter conducted similarly as in the *3OW* (combining four comparison maps instead of two). Similarly, as for *3OW* red and yellow denotes clusters from the above-mentioned analyses, while red and blue from the same analysis, where all object category level (i.e. voxel pattern across all exemplars) has been controlled for in the regression analysis I.e. the combination analysis shows for each object category the regions where the attended object voxel pattern significantly better predicted the voxel pattern of the scene than the two distractors and when the object was a distractor itself.

As can be seen in Fig 7D-H, the way attention modulated spatial activation patterns depended strongly on both experiment (*3OW* and *3OW)* and attended object (*human, instrument, animal*). For speech, in *3OA* the attended exemplar dominated voxel patterns in extensive bilateral AC fields, temporal cortex as well as left inferior–lateral frontal regions (Fig 7 D, yellow and red). However, when controlling for the average speech related activation pattern only left lateral frontal regions remained (Fig 7D, blue and red). In contrast, in *3OW* the attended speech exemplar dominated voxel patterns in right AC fields (TA2, STGa, STSda, PI), as well as right middle temporal gyrus, right insular fields, right temporal pole and bilateral inferior parietal–sensorimotor regions (Fig 7C, yellow and red). When controlling for the average speech related activation pattern the exemplar related pattern in the right AC was skewed posteriorly to subfields PBlelt, A4, and A5 (Fig 7C blue and red).

For animals, in *3OA* the attended exemplar dominated activation patterns at the border between the insula, PI and 52, irrespective of whether the average animal related activation pattern was controlled for or not (Fig 7F, red). In contrast, right motor and inferior frontal regions only emerged when controlling for the average animal pattern (Fig 7F, blue). In *3OW*, the attended animal exemplar dominated the same right AC subfields as observed for speech in *3OW* and additionally the subfields MBelt and PBelt (Fig 7E, red and yellow). Controlling for the average animal spatial pattern had little effect on the results in AC for animal exemplars in *3OW* (Fig 7E, red). Outside the AC attended animal exemplars dominated voxel patterns in bilateral middle temporal gyrus extending to the temporal pole, as well as bilateral inferior parietal–sensorimotor regions and the right insula extending to inferior frontal cortex (Fig 7, red and yellow).

For instruments, in *3OA* the attended instrument exemplar dominated activation patterns bilaterally in AC fields Ta2 and PI. Outside the AC the attended instrument exemplars dominated spatial patterns in bilateral inferior frontal, bilateral insular, left lateral frontal, and right middle temporal gyrus extending to the temporal pole (Fig 7H, red and yellow). Controlling for average instrument spatial patterns had little effect on the results except for the right AC. In *3OW*, the attended instrument exemplar dominated similar bilateral AC spatial activation patterns as in *3OA* (albeit right AC subfields only emerged when controlling for average instrument activation patterns, Fig 7G red and yellow). Outside the AC attended instrument exemplars dominated spatial activation patterns in bilateral insula, lateral visual cortex, left temporal pole, left inferior frontal, and sensorimotor regions.

## Discussion

We investigated how selective attention modulates neural representations of distinct categories of sounds (*speech, instrument, animal*) in auditory scenes comprising three overlapping sound objects. The three different classes of sound objects activated distinct subfields of the AC irrespective of attention. Speech objects were associated with higher-level lateral AC subfields, while animal and instrument objects were associated with higher-level medial AC subfields. As expected, selective attention modulated sound processing in the respective object type selective AC fields. This supports the notion that selective attention increases the gain of neurons that encode the stimulus features of the sounds through top-down neural signals propagating from higher-level cortical regions (Kerlin et al., 2010; Zion-Golumbic et al., 2013). However, our spatial pattern analysis results showed that this is only one of the potential neural mechanisms by which selective attention transforms neuronal representations in AC. We found that selective attention influences auditory representations in a highly contextual manner, depending on both the attended object type and the sound scene in which the object is embedded.

### Stimulus dependent selectivity for speech, instrument and animal auditory objects in AC subfields

The lateral subfields of AC (A4, A5), as well as STG/STS subfields (STGa, STSda, STSdp) showed selectivity for speech objects, while medial AC subfields (PI and 52) showed selectivity for instrument and animal objects, corroborating models where processing of speech diverges from other auditory objects in higher-level fields of AC (Hickok, 2010; Zhang et al., 2011; Friederici and Gierhan, 2013; Rolls et al., 2023). The exact location of the primary and secondary AC subfields in humans is highly debated (Leaver and Rauschecker, 2016). Nevertheless, our object selective subfields fall outside of tonotopically organized fields of the human AC (Formisano et al., 2003; Langers et al., 2007; Saenz et al., 2011; Moerel et al., 2014; Leaver and Rauschecker, 2016) and fields selective for standard acoustic dimensions (Leaver and Rauschecker, 2010b; Norman-Haignere et al., 2015; Hausfeld et al., 2018; Erb et al., 2019; Giordano et al., 2023). Our finding that these AC subfields support separate functions in object perception is also corroborated by a recent functional connectivity study of human AC. This study showed that the medial and lateral AC subfields are highly interconnected and connected to putative primary and secondary AC fields, but connections between the medial and lateral subfields are sparse (Rolls et al., 2023). Thus, these object selective subfields may correspond to parabelt fields in the macaque AC (Rauschecker and Tian, 2000). Note, however, that medial parabelt fields have been poorly characterized in macaques (Kusmierek and Rauschecker, 2009).

Our speech selective lateral cluster on the one hand and medial animal/instrument cluster on the other (Fig 4) coincides with AC clusters that have been identified as categorically speech vs. music selective (Norman-Haignere et al., 2015; Hausfeld et al., 2018; Norman-Haignere et al., 2022). Instead of using identifiable music, our instrument sounds were generated from human speech by extracting the pitch profile of the speech sounds (see *Stimuli*). Thus, unlike speech vs. music, our speech and instrument sound objects were similar in both their low-level acoustic features (see Fig 6) and temporal structure (rhythm and predictability). Therefore, we argue that the selectivity of these fields does not seem to stem from a difference in processing speech and music, per se, but rather human speech and instrumental sound objects. The speech and instrument sound objects did differ in their timbral structure, which has been found to involve distinct processing in AC (Deike et al., 2004; Bizley et al., 2009; Bizley and Walker, 2010; Bizley et al., 2013). We, however, do not believe that the object selectivity observed in the medial AC relates to timbral selectivity, per se, as fields were not only selective for instruments vs. speech but also for animal sounds vs. speech. As there is little prior data on the function of these medial AC fields, it is difficult to define the exact role of these medial subfields, other than that they may process higher-level non-speech object properties or possibly integrate low-level feature information that differentiate such objects (Rauschecker and Scott, 2009).

We also found stronger stimulus dependent processing for instrument sounds than animal sounds in MBelt, LBelt and TA2 (Fig 3C, blue and turquoise). Similar sound object selectivity has not, to our knowledge, been observed in previous studies. However, a previous fMRI study (Norman-Haignere et al., 2015) that used naturalistic sound clips and inferred canonical response profiles (“components”) to explain voxel responses in AC, found that the sounds that elicited the strongest responses in these subfields all contained pitch or spectral modulations, reflecting harmonics. Thus, a difference in harmonic structure between animal and instrument sounds may explain the observed selectivity for instrument sounds above animal sounds in these subfields see also (Hausfeld et al., 2018).

### Attention modulates auditory processing in auditory object selective AC subfields

Attention related modulation (*ARM*) of sound scenes was found to show comparable object selectivity as stimulus dependent processing in AC. That is, medial AC subfields were more strongly activated when participants attended to animal/instrument than speech objects, while lateral AC subfields showed the reverse effect. This pattern of results was most evident in the *3OA-experiment*, where the sound scenes comprised sounds from separate auditory categories (Fig 4A-C). However, our Bayesian ROI analyses showed that similar object selectivity was also observed in the *3OW-experiment* albeit at a subthreshold level. A long line of research has found that selective attention to speech in sound scenes with several overlapping speakers is associated with activations in lateral AC subfields and STS regions as well as inferior frontal fields (e.g., Alho et al., 2003; Alho et al., 2006; Leminen et al., 2020a; Wikman et al., 2020; Wikman et al., 2022; Ylinen et al., 2022). Our results, however, expand upon this literature by showing that these attentional modulations are speech specific, and do not reflect general attentional modulation of sound representations in AC.

In contrast to speech, instrument and animal related *ARMs* have been poorly characterized in previous studies. However, as for speech, *ARMs* for instrument and animal sounds generally coincided in the medial AC subfields (PI and 52) that also showed stimulus related sound object selectivity for these two categories. Many models of auditory object perception suggest that attention is necessary in forming stable object percepts acting as “glue” that integrates disparate object features in AC (Shinn-Cunningham, 2008; Shamma et al., 2010; Bizley and Cohen, 2013; Shinn-Cunningham et al., 2017). Thus, it was somewhat unexpected that within AC only two subfields (see Fig 4C red, yellow) showed both object selective stimulus dependent processing and object selective *ARMs*. Thus, either attention is not necessary for object formation; attention mediated object formation occurs outside AC subfields (e.g., inferior frontal cortex (Romanski, 2012)); or attention facilitates sound object formation in the same subfields that encode features that differentiate the different types of sound objects. It is also possible that the neuronal populations representing whole auditory objects or object classes are so sparse that the resolution of fMRI is not sufficient (Leaver and Rauschecker, 2010b).

Object specific ARMs generally occurred in subfields of AC that have not been found to represent low-level acoustic features. However, since it has been shown that selective attention can be tuned to low-level features (Shamma et al., 2011; Da Costa et al., 2013), we determined whether any of the categorical differences in ARMs could be explained by the acoustic features of the attended object in our *3OA-experiment*. Based on a prior study (Leaver and Rauschecker, 2010a), we chose spectral (*pitch* and *frequency centroid*), spectral structure (*harmonic to noise ratio*), and temporal variability (*frequency modulation standard deviation* and *amplitude modulation standard deviation*) features. In addition to these acoustic features, we investigated whether *entropy*, which describes the overall information level of the sound clip affected ARMs.

Unlike prior studies which have included all acoustic features and categorical variables in the same *GLM* we chose to run each analysis separately. This was chosen because of strong multicollinearity and negative correlations between the features, which would have yielded possibly uninterpretable results. As can be seen in Fig 6 several of the acoustic features correlated with ARMs in object selective subfields of AC. However, all these correlations seemed to arise because speech sound objects differed from the two other object categories on these acoustic features (Fig 6A, C, D). In contrast, ARMs the medial subfields PI and 52, where both stimulus and attention related selectivity for animal and instrument sound objects was observed, did not correlate with any of the studied low-level features. Instead, we observed that ARMs in LBelt, PBlelt and MBelt showed positive correlations with *pitch* and *frequency centroid* and *entropy.* These patterns bear a striking resemblance to the high-frequency selective tonotopic AC subfields observed in the study of (Norman-Haignere et al., 2015) and probably relate to the processing of high frequency components in the sound scenes.

We want to highlight that although no single low-level acoustic feature *alone* seemed to explain the object selectivity observed for *ARMs* in AC, we do not claim that our data suggest that attention *only* modulated higher-level object processing and was not tuned based on lower-level features such as frequency at all. Whether attention can be tuned to low-level spectrotemporal features is a relatively pertinent question as several previous fMRI e.g., (Alho et al., 1999; Petkov et al., 2004; Rinne et al., 2009) and EEG studies e.g., (Ding and Simon, 2012; Akram et al., 2016) studies have argued that attention modulates sound processing only at late stages of cortical processing, i.e., after initial spectrotemporal analysis. Accordingly, we observed most of the *ARM* differences in AC subfields that have not been associated with spectrotemporal response properties (see *Stimulus dependent selectivity for speech, instrument and animal auditory objects in AC subfields*). However, we also for example observed that attending to speech in *3OA* caused stronger *ARMs* in A1 than attending to animals in similar scenes (Fig 4A, see also differences between ARMs to instruments and those to animals Fig 4C). Although the HCP-parcellations A1 might not correspond exactly to the primary AC, the field lies in the center of tonotopically organized regions (Norman-Haignere et al., 2015), with spectral tuning for low frequency. Thus, it is initially perplexing that this field would show differences between animal and speech attention. However, because speech and animal objects differed in their spectral decomposition (Fig 5), with human sound objects on average including more low-frequency components than the animal objects, attentional precedence for the speech objects could be given by boosting sound processing in low-frequency spectrotemporal fields of the AC. Thus, rather than arguing that attention cannot influence lower-level sound processing, per se, we suggest that attention adaptively modulates the level of sound processing that offers the necessary contrast between the attended and distractor sounds (see also *Attentional selection of auditory objects depends on the auditory scene*).

### Attentional selection of auditory objects depends on the auditory scene

Attention is classically assumed to enhance the gain or tuning properties of feature selective neurons in AC (Fritz et al., 2007; Kauramäki et al., 2007; Moerel et al., 2013). This gain increase has also been suggested to explain why neurons entrain to the changes in task relevant sounds at the cost of task irrelevant sounds (Kerlin et al., 2010; Mesgarani and Chang, 2012; Power et al., 2012; Zion-Golumbic et al., 2013; O’Sullivan et al., 2015; O’Sullivan et al., 2019). Our findings that attention selectively modulate processing in the same AC subfields responding to stimulus processing of the attended sound object is in line, but by no means proves this notion. Thus, to test whether this model explains attentional dynamics in the current data we performed a spatial pattern analysis. We conjectured that if attention simply increases the gain of the neurons processing attended objects, then the spatial activation pattern of the auditory scenes should be dominated by the attended auditory object exemplar, at the cost of the distractor exemplars, in respective auditory object selective AC fields (Hausfeld et al., 2018; Puschmann et al., 2024). Further, if attention only influences neuronal gain in AC, this attentional effect should not depend on whether the distractors are comprised of objects from the same object category (*3OW*) or objects from different object categories (*3OA*). As can be discerned in Figure 7, how attention affected spatial activation patterns strongly depended on both the sound scene type (*3OA vs. 3OW*) and attended object (*speech, instrument, animal*) suggesting that attentional dynamics are much more complex than simple neuronal gain increase, as follows.

For speech, we found that the attended speech object exemplar dominated the spatial activation pattern of the sound scene across the whole left hemisphere language processing network (Friederici and Gierhan, 2013; Fedorenko et al., 2024). However, this applied only to *3OA* where each object of the scene comprised a different category. The same analysis in *3OW,* where both the attended object and the distractors consisted of speech, yielded strikingly dissimilar results. In *3OW* the attended speech dominated activation patterns in the right hemisphere rather than the left, and in AC the cluster was centered on fields representing low-level speech properties (Norman-Haignere et al., 2015; Norman-Haignere et al., 2022) extending to AC fields responsive for low-level sound features (Norman-Haignere et al., 2015). When general speech category level spatial activation patterns (across all speech object exemplars) were controlled for, the extensive language processing network correlates did not remain significant in *3OA,* while the effect of average speech related spatial patterns was much more nuanced in *3OW*. Thus, it seems that in *3OA* the general speech related activation pattern was boosted by attention, while in *3OW* the attentional dynamics were entirely different. Importantly, differences between the two experiments in attentional processing of speech were not only confined to our spatial pattern analysis but also our univariate analysis of ARMs indicated stronger ARMs in *3OW* than in *3OA* in posterior auditory fields (A5, STSda, STSdp, Fig 6A). The disparity between attentional dynamics in the two experiments was not fully unprecedented, as we conjectured that in *3OW* attention might operate on lower-level features to separate objects within category (*3OW*), while category-level differences could be used in *3OA* to separate the three objects. Accordingly, the right hemisphere AC subfields that showed attentionally enhanced spatial patterns in *3OW* included fields that are responsive to lower-level spectral and temporal modulation (Norman-Haignere et al., 2015). Together this may suggest that the neural networks that differentiate speech from other categories of sounds are distinct and only partially overlapping with those that differentiate between different speakers (Giordano et al., 2013), with attention operating on different levels of representation when necessary (Wikman et al., 2020; Wikman et al., 2024b). Further, our results also support previous findings where right hemisphere dominance has been observed for speaker identity differentiation which was only necessary in *3OW* and not 3OA where there was only one speaker in the scene (Bonte et al., 2014).

In *3OA,* only the instrument/animal selective medial AC cluster (see Fig7E) showed stronger correlates with attended animal exemplars than the distractors, while in *3OW* correlates in AC where right lateralised and comprised both subfields selective for objects and subfields responsive to spectral and temporal modulation. Thus, as for speech objects, the results are consistent with the notion that attention boosts different aspects of auditory objects in scenes comprised of within category exemplars and between category exemplars. In contrast, attentional spatial pattern effects for instruments were more comparable in the two experiments (*3OA vs. 3OW*), with attentional precedence occuring in the same AC subfields that selectively responded to intrument object features (see Fig 7G–H) and AC subfields responsive to spectral modulation of sounds. The latter being especially evident when controlling for general instrument related activation patterns. Importantly, unlike speech, controlling for general object category level spatial patterns did not affect the spatial pattern analysis results strongly in AC for either animal or instrument exemplars. This may indicate that each animal and instrument exemplar is represented by its idiosyncratic pattern in AC, albeit representations overlap in object responsive medial AC subfields. This notion is consistent with the view that supra category level representations of natural objects are not present at the level of the auditory cortex (Bizley and Cohen, 2013).

Our result that attended speech objects dominated right hemisphere AC spatial activation patterns instead of left hemispheric speech sensitive auditory fields in scenes comprised overlapping speech is in stark contrast to a recent study by (Puschmann et al., 2024). In this study participants attended to one of two overlapping speech streams, i.e. “cocktail party speech” during fMRI. Similarly to the current study, the authors used the spatial activation pattern of the attended speech stream and the distracting speech stream, each presented in isolation, to predict the spatial activation pattern of the overlapping speech streams. In contrast to our results, however, the authors found that the attended input predicted spatial activation patterns in higher level bilateral AC and STS regions more strongly than the distracting speech input.

One important difference between our and this previous study may explain the conflicting results. The study of Puschmann used long speech streams (> 30 mins), while we used only short speech segments (< 10 s). Further, Puschmann restricted their analysis to the end of the timeseries, i.e., did not analyse the first minute of the overlapping speech streams. We argue the disparity between our results and those of Puschmann might be explained by two distinct and temporally partially overlapping roles of attention in speech sound processing. First, as discussed above, attention affects stream segregation in AC (Griffiths and Warren, 2002; Griffiths and Warren, 2004; Angeloni and Geffen, 2018) by “boosting” the neural processes that differentiate attended objects from distracting sounds. We have previously argued that this relates to the building a so-called “attentional map” (Näätänen, 1990) in AC representing both low- and higher-level properties that differentiate attended objects from distractors (Wikman et al., 2020; Wikman et al., 2024b). We argue that when this map is stable a second attention driven phenomenon occurs, namely attention starts to gate higher-level information prosessing (semantic, lingusitic etc.) preferentially for the attended speech input and supress such information processing for distracting speech input (Puschmann et al., 2024). Thus, as our analysis was restricted to less than 10 seconds of overlapping speech, attention was not yet gating exclusively the attended input for higher-level processing. In contrast, in the study of Puschman et al, the characteristics differentiating the two streams had probably already been established. This model is compatible with previous results indicating that semantics of distracting speech is prossessed only in the beginning of sentences (Näätänen et al., 1992; Wikman et al., 2024a), when speech streaming is incomplete. Further this offers an attractive account for some of the striking dissimilaities observed between speech attention effects on spatial activation patterns in *3OA* and *3OW.* That is, when the distracting auditory input is non-speech (*3OW*) the characteristics differentiating speech from the distractor sounds are immediately evident and the attentional map might be formed almost instantaneously and thus higher level analysis of speech can be much more quickly gated by attention, explaining why correlates where observed across the language network. Thus, our study offers tantalising new evidence that temporal aspects of attentional entrainment are important to consider, i.e., selective attention oprates on several levels ad different points in time, in a context and task dependent manner (Wikman et al., 2022; Wikman et al., 2024b).

Lastly, it is evident that the spatial analysis results were strikingly dissimilar between speech and animal/instrument sound objects, indicating poor generalisation of attentional effects between speech and other types of auditory objects. Future studies using incomprehensible speech or non-speech human vocal sounds are, however, needed to discern if this entirely relates to the precence of semantics and other linguistic properties in speech (Wikman et al., 2022; Fedorenko et al., 2024; Wikman et al., 2024b), or whether attentional entrainement to human vocalisations in general utilse distinct neural mechanisms.

### Attentional effects outside AC subfields

This paper focuses on attentional dynamics of neural auditory scene representation in AC. However, we want to highlight some observations on attentional dynamics outside AC. Firstly, we replicated that selectively attending to speech is associated with stronger left hemisphere inferior frontal regions (Wikman et al., 2020; Wikman et al., 2022; Ylinen et al., 2022; Wikman et al., 2024b) and regions associated with speech semantics (Binder et al., 2009). Further, attentional effects for speech in inferior frontal regions were stronger in *3OW* than in *3OA,* supporting models where inferior frontal regions act as primary control networks for speech attention, showing a graded response according to the demands of speech stream segregation (Wikman et al., 2020). Athough attentional effects in the inferior frontal cortex were stronger for speech in the left hemisphere (Fig 4), our Bayesian analysis indicated that bilateral inferior frontal regions also showed stronger activations in *3OW* than *3OA* when attending to animal and instrument sound objects. Thus, it may be that inferior frontal regions have a more general role in controlling attentional resources, showing limited specificity for speech attention. Lateral frontal regions, in contrast, were associated with attention to the non-speech objects rather than speech object. This finding is consistent with models where the level of frontal involvement in different tasks depends on task automaticity, with highly automatized tasks, such as listening to speech in noise, recruiting mainly sensory networks, while novel tasks (listening to animals or instruments in complex scenes) recruit the prefrontal cortex (Chein and Schneider, 2012). Lastly, we want to highlight that attentional dominance of spatial activation patterns did not merely occur in auditory and language network regions, but consistently also in the middle temporal gyrus extending to the anterior temporal lobe. This finding was especially evident in the *3OW* experiment and may indicate the importance of these cortical regions for categorical perception, possibly supporting formation of supracategorical representations (Lewis et al., 2011; Theunissen and Elie, 2014).

## Notes

### Competing Interest Statement

The authors have declared no competing interest.

